# Single-Cell Mass Cytometry on Peripheral Blood Identifies Immune Cell Subsets Associated with Primary Biliary Cholangitis

**DOI:** 10.1101/2020.02.24.962043

**Authors:** Jin Sung Jang, Brian Juran, Kevin Y. Cunningham, Vinod K. Gupta, YoungMin Son, Ju Dong Yang, Ahmad H. Ali, Elizabeth Ann L. Enninga, Jaeyun Sung, Konstantinos N. Lazaridis

**Affiliations:** Medical Genome Facility, Center for Individualized Medicine, Mayo Clinic, Rochester, MN, USA; Department of Laboratory Medicine and Pathology, Mayo Clinic, Rochester, MN, USA; Division of Gastroenterology and Hepatology, Department of Internal Medicine, Mayo Clinic, Rochester, MN, USA; Graduate Research Education Program (GREP), Mayo Clinic, Rochester, MN, USA; Department of Computer Science and Engineering, University of Minnesota Twin-Cities, Minneapolis, MN, USA; Microbiome Program, Center for Individualized Medicine, Mayo Clinic, Rochester, MN, USA; Division of Surgical Research, Department of Surgery, Mayo Clinic, Rochester, MN, USA; Division of Pulmonary and Critical Care Medicine, Department of Medicine, Mayo Clinic, MN, USA; Department of Immunology, Mayo Clinic, Rochester, MN, USA; Division of Digestive and Liver Diseases, Department of Medicine, Cedars Sinai Medical Center, Los Angeles, CA, USA; Department of Obstetrics and Gynecology, Mayo Clinic, Rochester, MN, USA; Division of Rheumatology, Department of Medicine, Mayo Clinic, Rochester, MN, USA

**Keywords:** Primary Biliary Cholangitis, Liver Cirrhosis, Mass Cytometry, Immunophenotyping, Cluster Analysis, Machine Learning

## Abstract

The relationship between Primary Biliary Cholangitis (PBC), a chronic cholestatic autoimmune liver disease, and the peripheral immune system remains to be fully understood. Herein, we performed the first mass cytometry (CyTOF)-based, immunophenotyping analysis of the peripheral immune system in PBC at single-cell resolution. CyTOF was performed on peripheral blood mononuclear cells (PBMCs) from PBC patients (n=33) and age-/sex-matched healthy controls (n=33) to obtain immune cell abundance and marker expression profiles. Hiearchical clustering methods were applied to identify immune cell types and subsets significantly associated with PBC. Subsets of gamma-delta T cells (CD3^+^TCRgd^+^), CD8^+^ T cells (CD3^+^CD8^+^CD161^+^PD1^+^), and memory B cells (CD3^-^CD19^+^CD20^+^CD24^+^CD27^+^) were found to have lower abundance in PBC than in control. In contrast, higher abundance of subsets of monocytes and naïve B cells were observed in PBC compared to control. Furthermore, several naïve B cell (CD3^-^CD19^+^CD20^+^CD24^-^CD27^-^) subsets were significantly higher in PBC patients with cirrhosis (indicative of late-stage disease) than in those without cirrhosis. Alternatively, subsets of CD8^+^CD161^+^ T cells and memory B cells were lower in abundance in cirrhotic relative to non-cirrhotic PBC patients. Future immunophenotyping investigations could lead to better understanding of PBC pathogenesis and progression, and also to the discovery of novel biomarkers and treatment strategies.

## Introduction

Primary Biliary Cholangitis (PBC) is an autoimmune liver disease characterized by immune infiltration and targeted destruction of intrahepatic bile ducts. This results in chronic cholestasis and ultimately in progression to cirrhosis and liver failure^1-3^. PBC predominantly affects women (90% of patients), and serum antimitochondrial antibodies (AMA) specific to the E2 subunits of 2-oxoacid dehydrogenase complexes are present in 90-95% of patients^4,5^. These autoantibodies provide bile duct specificity due to a unique aspect of biliary epithelial cell apoptosis, which leaves the E2 subunit of the pyruvate dehydrogenase complex immunologically intact and available on apoptotic blebs^6^.

Recent insights suggest that the offending immunological processes may vary depending on disease stage/activity, and that individual differences in immune response are likely to contribute to variability in disease course^3^. As such, personalized approaches to treatment would be highly beneficial; however, direct and routine assessment of immune activity in the liver is precluded due to the invasiveness and risks associated with liver biopsy. Moreover, biopsy material is quite localized and not always reflective of disease state throughout the whole liver. A number of studies focused on peripheral immunity in PBC have reported alterations in various immunological subsets including natural killer (NK) cells^7^, regulatory T cells^8^, CD8^+^CD57^+^ T cells^9^ and mucosal-associated invariant T (MAIT) cells^10^. To what extent these differences reflect ongoing liver pathology, or directly implicate pathological subsets, remains to be determined. However, the use of peripheral blood to assess alterations to immune composition allows for the incorporation of much larger sample sizes into studies, and facilitates longitudinal observations that may ultimately prove clinically relevant.

Mass cytometry is an emerging platform for immunophenotyping that overcomes key limitations of traditional fluorescence-based flow cytometry^11,12^. This technology utilizes heavy-metal ion tags instead of fluorophores to detect target-bound monoclonal antibodies, allowing for simultaneous quantification of 40 or more cell markers^11^. This increased dimensionality facilitates a more comprehensive survey of immune composition and has recently been employed in the examination of peripheral blood in various pathologies, including autoimmune diseases and cancers^12–14^.

In this pilot study, we aimed to evaluate the use of mass cytometry to immunophenotype stored peripheral blood mononuclear cells (PBMCs) collected from PBC patients and healthy subjects (controls) in order to study differences in immune cell subsets, as well as to inform future studies utilizing our extensive biobank blood collections.

## Methods

### Study population

All participants in this study, i.e., well-documented PBC patients and clinic-based controls, were previously recruited into our Mayo Clinic PBC Genetic Epidemiology (MCPGE) Registry and Biospecimen Repository, which was initiated with the aim to elucidate the genetic and environmental contributors to PBC pathogenesis^15^. Diagnosis of PBC was based on the published American Association for the Study of Liver Diseases (AASLD) criteria^16^. Cirrhosis was diagnosed by liver biopsy, radiology, and/or portal hypertension documented by the presence of ascites and/or subsets. Patient selection included only female patients who tested AMA positive and were taking ursodeoxycholic acid (UCDA) at the time of sample collection. Controls were individually matched to patients based on sex, age at sample collection (+/- 1 year) and date of sample collection (+/- 1 year). Demographic and clinical characteristics of participants are provided in **Table 1**. All blood samples were obtained from study participants following written informed consent. This study was approved by the Mayo Clinic Institutional Review Board in accordance with the Declaration of Helsinki. All methods and procedures were performed in accordance with Mayo Clinic Institutional Review Board guidelines and regulations.

**Table 1.** General characteristics of the study subjects.

| Characteristics | PBC (range) | Control (range) |
| --- | --- | --- |
| Number of subjects | 33 | 33 |
| Age at initial diagnosis (years) | 50.6 (37–63) | N/A |
| Age at sample collection (years) | 57.9 (50.1–64.8) | 57.9 (50.4–64.8) |
| Duration between diagnosis and sample collection (years) | 7.28 (0.0–23.4) | N/A |
| Gender (female) | 33 (100%) | 33 (100%) |
| Race (caucasian) | 33 (100%) | 33 (100%) |
| Non-cirrhosis/cirrhosis/unavailable | 26/6/1 | 33/0/0 |
| Alkaline phosphatase (U/L)<sup>†</sup> | 185.7 ± 131.2 | 78.0 ± 19.5 |
| AMA (positive/negative) | 33/0 | 0/33 |
| UDCA treatment (yes/no) | 33/0 | 0/33 |

<sup>†</sup>mean ± standard deviation. N/A, not applicable. AMA, serum antimitochondrial antibodies. UDCA, ursodeoxycholic acid.

### Cell isolation, preparation, and labeling

Human PBMCs were isolated using Ficoll-Paque density-gradient centrifugation (GE Healthcare, NJ), slow-frozen and stored in liquid nitrogen until preparation for mass cytometry. Frozen PBMCs were thawed at 37°C, combined with 1 mL of cell media (RPMI, 10% FBS, Pen/Strep), centrifuged at 1500 RPM for 5 min and resuspended in 1 mL of warm cell media. Cells were then counted on a Countess II automated cell counter and approximately 3×10^6^ cells (in 1 mL volume) of each PBMC sample was prepared and incubated at 37°C for 1 hr prior to labeling. Cell labeling was performed as per manufacturer recommendations (Fluidigm Sciences). Briefly, cells were isolated, resuspended in 0.5 µM Cell-ID cisplatin solution (Fluidigm Sciences) and incubated at room temperature for 5 min to stain dead cells. Cells were then washed twice with 1 mL Maxpar Cell Staining Buffer (MCSB, Fluidigm Sciences) and resuspended in 50 µL of MCSB. To this, 50 µL of antibody cocktail consisting of 36 metal-conjugated antibodies in MCSB was added and samples were incubated at room temperature for 45 min with gentle agitation. The antibodies were obtained from Fluidigm or generated in-house by the Mayo Clinic Hybridoma Core using Maxpar X8 Ab labeling kits (Fluidigm) and are detailed in **Supplementary Table S1**. Following staining, cells were washed twice with 1 mL MCSB, resuspended in 1 mL of fixation solution (1.6% PFA in CyPBS) and incubated at room temperature for 20 min with gentle agitation. Fixed cells were then rinsed twice with 1 mL MCSB and cell pellets were stored overnight at 4°C. Pellets were next resuspended in 1 mL intercalation solution [62.5 nM Cell-ID Intercalator-Ir (Fluidigm Sciences) in MaxPar Fix and Perm Buffer (Fluidigm Sciences)] to which 50 µl of diluted barcoding solution prepared using the Cell-ID 20-Plex Pd Barcoding Kit (Fluidigm Sciences) was added and samples were incubated overnight at 4°C. Barcoded samples were washed with 1 mL MCSB, resuspended in 1 mL CyPBS and cells were counted on a Countess II automated cell counter. Finally, cells were resuspended in Cell Acquisition Solution-EQ Bead mixture (Fluidigm Sciences) to a concentration of 5×10^5^ cells/mL before loading onto a Helios CyTOF system (Fluidigm, CA).

### Mass cytometry and data acquisition

The barcoded samples were loaded onto a Helios CyTOF system using an attached autosampler and were acquired at a rate of 200–400 events per sec. Data were collected as .FCS files using CyTOF software (Version 6.7.1014, Fluidigm). After acquisition, intrafile signal drift was normalized to the acquired calibration bead signal and individual files were deconvoluted and stored into .fcs files using CyTOF software. File clean-up (e.g., removal of dead cells, debris, doublets, and beads) was performed using Gemstone software (Verity Software House).

### Identification of immune cell subsets associated with PBC using clustering analyses

Gemstone-cleaned .fcs files were used for subsequent analyses in the Cytobank cloud-based platform (Cytobank, Inc.). First, all 66 files (corresponding to the 66 study subjects) were uploaded onto Cytobank. viSNE, which is a dimensionality reduction technique for high-dimensional single-cell data based upon the Barnes-Hut implementation of t-SNE^17^, was then used to visualize the mass cytometry data as 2D t-SNE maps with the following parameters: Desired Total Events (with Equal Sampling): “100,000” (per group); Channels: select all 36 antibody-metal channels; Compensation: “File-Internal Compensation”; Iterations: “2000”; Perplexity: “60”, and Theta: “0.75”. For more advanced data visualization and exploration of cytometry data to identify clinically-relevant immune cell subsets, two additional computational methods, FlowSOM and CITRUS, were used with default settings unless otherwise noted. FlowSOM uses Self-Organizing Maps (SOMs) to partition all individual cells based on their marker expression phenotypes into clusters and metaclusters (i.e., groups of clusters) and provides their global connections in the format of a Minimum Spanning Tree (MST)^18^. Following the generation of an MST, each cluster and metacluster can be queried for its immune cell abundance and distribution of different cell surface markers. FlowSOM was performed with the following parameters: Event Sampling Method: “Equal”; Desired events per file: “17,178”; Total events actually sampled: “1,133,748”; SOM Creation: “Create a new SOM”; Clustering Method: “Hierarchical Consensus”; Number metaclusters: “20”; Number clusters: “256”; Iterations: “10”; Seed: “1234”. CITRUS (cluster identification, characterization, and regression) identifies clusters of cell subpopulations that are statistically associated or correlated with an experimental or clinical phenotype of interest (e.g., disease/control, responder/non-responder)^19^. The output is a network topology of cell subpopulation clusters that represents a hierarchical stratification of the original samples. Features that drive the differentiation between phenotypes may be either the relative abundance of cell subpopulations or the median expression levels of functional markers measured across cells of each population. CITRUS was performed using the Significance Analysis of Microarrays (SAM) correlative association model [Benjamin-Hochberg-corrected *P*-value, i.e., false discovery rate (FDR)<0.01] with the following parameters: Clustering channels: “select all 36 antibody-metal channels”; Compensation: “File-Internal Compensation”; Association Models: “Significance Analysis of Microarrays (SAM) - Correlative”; Cluster Characterization: “Abundance”; Event sampling: “Equal”; Events sampled per file: “5,000”; Minimum cluster size (%): “1”; Cross Validation Folds: “5”; False Discovery Rate (%): “1”. Identification of differentially abundant immune cell subsets between groups was performed by comparing the mean of the relative abundances of each FlowSOM cluster, of each FlowSOM metacluster, or of each CITRUS cell subpopulation cluster. Unless otherwise noted, the Mann-Whitney *U* test was used for all statistical hypothesis testing, with a minimum fold-change of 2 and *P*-value < 0.05 required to be considered as statistically significant.

### Training a neural network with mass cytometry data features for supervised classification

The Python version of the H2O AutoML package (version 3.26.0.3) was used to train a neural network model with stochastic gradient descent for distinguishing mass cytometry profiles from PBC and control subjects. On the training data, AutoML runs in parallel a large selection of candidate machine-learning algorithms with uniquely tuned parameters (using random grid-search) and provides the top-performing, parameter-tuned model for future cases. To run AutoML, H2O provides individual cloud instances utilizing Amazon Web Services (AWS). The input parameters that were used prior to model-training are the following: Epochs=“10,000”; Mini batch size=“1”; Random seed=“1234”. Data curation and model implementation was performed in Python version 3.6.4.

## Results

A schematic overview of our analysis of the PBMC immunophenotyping data is shown in **Fig. 1.** Briefly, PBMCs were collected from 33 female PBC patients and 33 age-/sex-matched controls, and subjected to single-cell mass cytometry (CyTOF) to obtain immune cell abundance profiles. A summary of the output data from our mass cytometry experiments, including total number of cells collected and stained for each sample, event counts detected by the instrument, and number of live/singlets for analysis, is provided in **Supplementary Fig. S1** and **Supplementary Table S2**. After post-processing of raw sample data obtained from the CyTOF machine, immune profiles were analyzed through the Cytobank cloud-based platform for the following: **i)** 2D-visualization and exploration of single-cell data using viSNE; and **ii)** identification of immune cell types and their subsets associated with PBC using hierarchical clustering approaches in FlowSOM and CITRUS.

**Figure 1.**
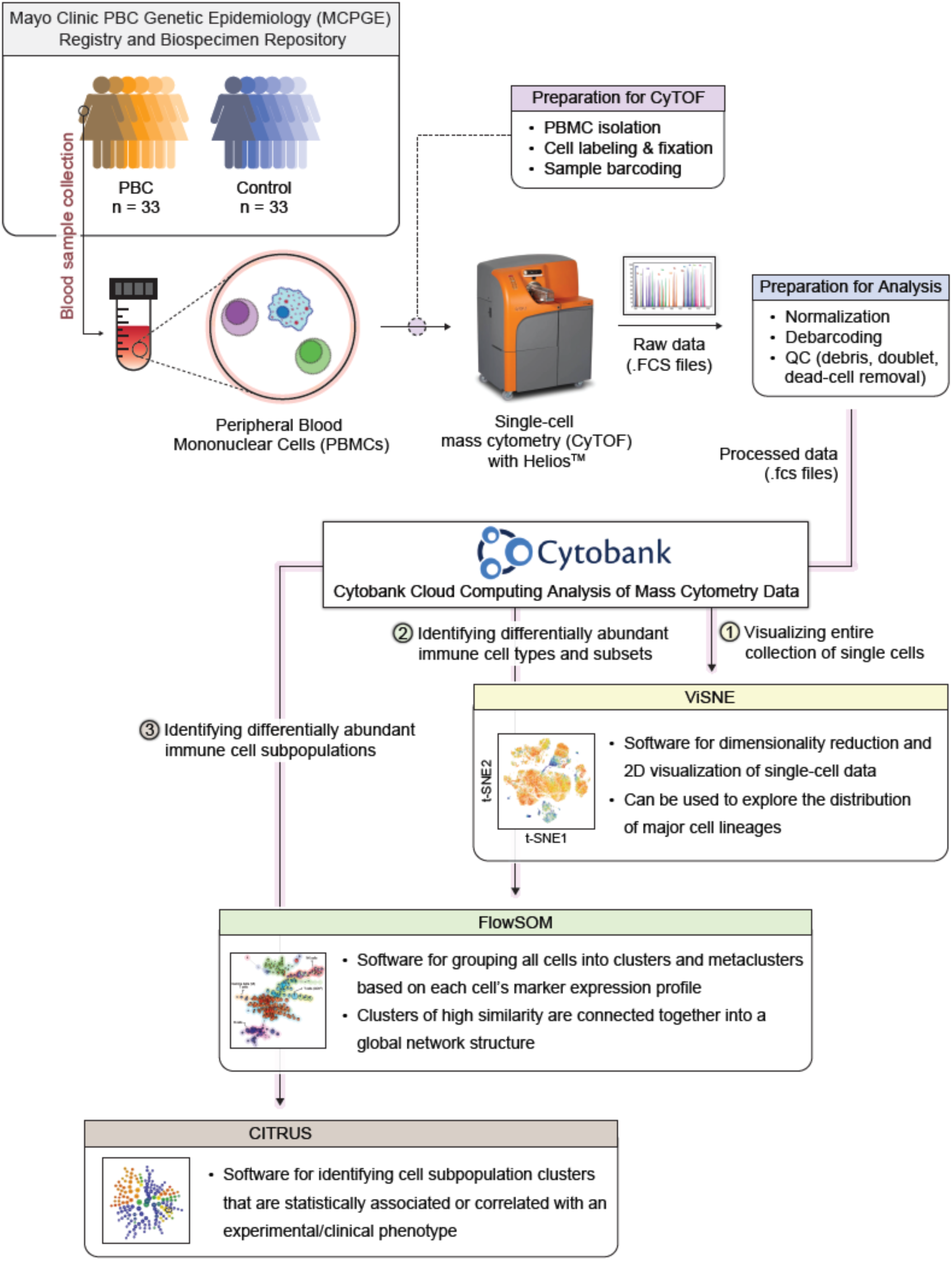
Schematic overview of the current study’s analysis pipeline to investigate immune cell populations implicated in PBC using the Cytobank cloud-based platform (Cytobank, Inc.).

### PBC patients and control subjects exhibit global differences in the peripheral immune system

To perform a qualitative analysis of the PBMC immunophenotyping data, we first applied viSNE on the immune cell abundance profiles from all PBC patients and control subjects. The resulting t-SNE maps for PBC and control groups show several distinct spatial regions (**Fig. 2A**). This finding reflects the heterogeneity of major immune cell lineages (e.g., T cells, B cells), which can be loosely defined using t-SNE maps colored by marker intensities (**Supplementary Fig. S2**), across immune profiles. Interestingly, we observed differences in densities of particular localized regions between the two maps, implying altered relative abundances of immune cell types and their subsets between PBC patients and controls. These regions of differential density can be broadly inferred to include B cells (CD3^-^, CD19^+^, CD20^+^; center far right), natural killer (NK) cells (CD3^-^, CD19^-^, CD14^-^, CD56^+^; far bottom center), CD4^+^ T cells (CD3^+^, CD4^+^; center near left), CD8^+^ T cells (CD3^+^, CD8^+^; upper left), and monocytes (CD3^-^, CD19^-^, CD14^+^; upper right). To further evaluate differences in immune cell distributions between PBC patients and controls, we used FlowSOM to generate cell clusters, which are visualized as a MST (**Fig. 2B**). As with viSNE, differences in B cell, T cell, NK cell and monocyte clusters between patients and controls are visually apparent.

**Figure 2.**
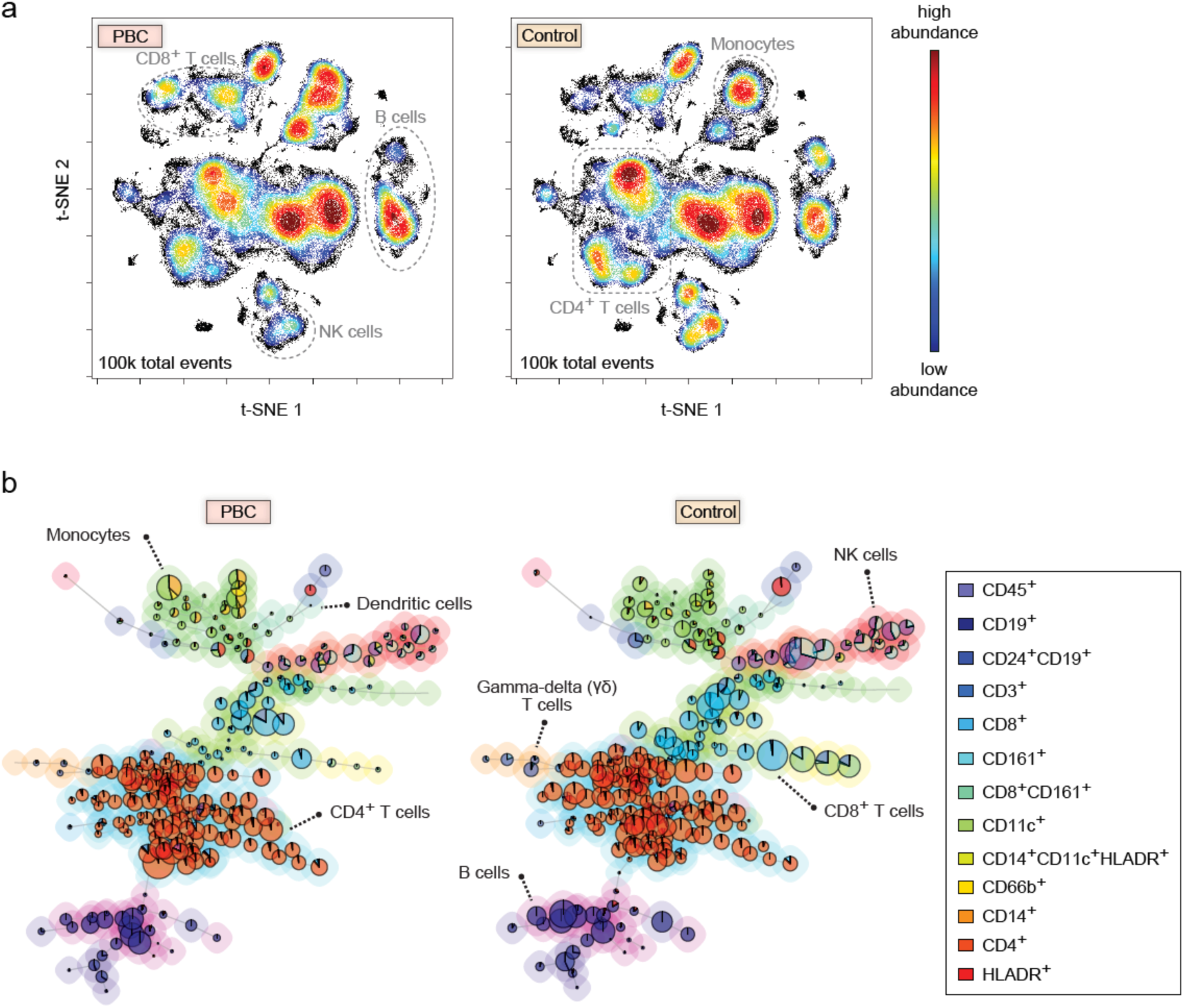
Qualitative analysis of PBMC immunophenotyping data using (A) viSNE and (B) FlowSOM reveals differences in immune cell lineages between PBC and control. (A) Mass cytometry samples from 33 PBC patients were concatenated into 100,000 randomly-sampled total events and mapped onto a t-SNE plot using viSNE (left). Analogously, samples from 33 age-/sex-matched controls were mapped using viSNE (right). Each point in the t-SNE plot represents a single event (e.g., cell) detected by the mass cytometer, and colors vary according to cell abundance density. Observed regional differences in cell densities correspond to differences in relative abundances of major immune cell lineages. (B) FlowSOM clusters cells into cell subsets based on their marker expression patterns, and generates a Minimum Spanning Tree (MST) of those clusters (left: PBC; right: control). Each node is characterized by a pie chart, whose diameter is proportional to the number of events, and whose colors indicate specific markers defined in the legend. The background colors group nodes into cell types that correspond to different major immune cell types (FlowSOM Metaclusters). Each link connects cell subsets of similar marker expression patterns. In the two MSTs of PBC and control, the nodes located in the same position correspond to the same cell subset.

### Major immune cell lineages differentially abundant between PBC and control

FlowSOM identified twenty metaclusters, of which three were found to be differentially abundant between the PBC and control groups (**Table 2** and **Figs. 3A–C**). The first, Metacluster-3 (**Fig. 3A**), expresses markers indicative of a gamma-delta T cell population (CD3^+^TCRgd^+^) and was 1.2-fold lower in abundance in PBC patients than in controls (0.9% vs. 1.1%; *p*<0.05). The second, Metacluster-4 (**Fig. 3B**) expresses markers consistent with a subset of CD8^+^ T cells that express CD161 and PD1 (CD3^+^CD8^+^CD161^+^PD1^+^) and was found to be 2.7-fold lower in abundance in PBC patients relative to controls (0.6% vs. 1.6%; *p*<0.001). Finally, Metacluster-16 (**Fig. 3C**) expresses markers consistent with a subset of memory B cells (CD3^-^ CD19^+^CD27^+^CD38^-^) and was 1.8-fold lower in abundance in PBC patients than in controls (1.7% vs. 3.1%; *p*<0.005).

**Table 2.**
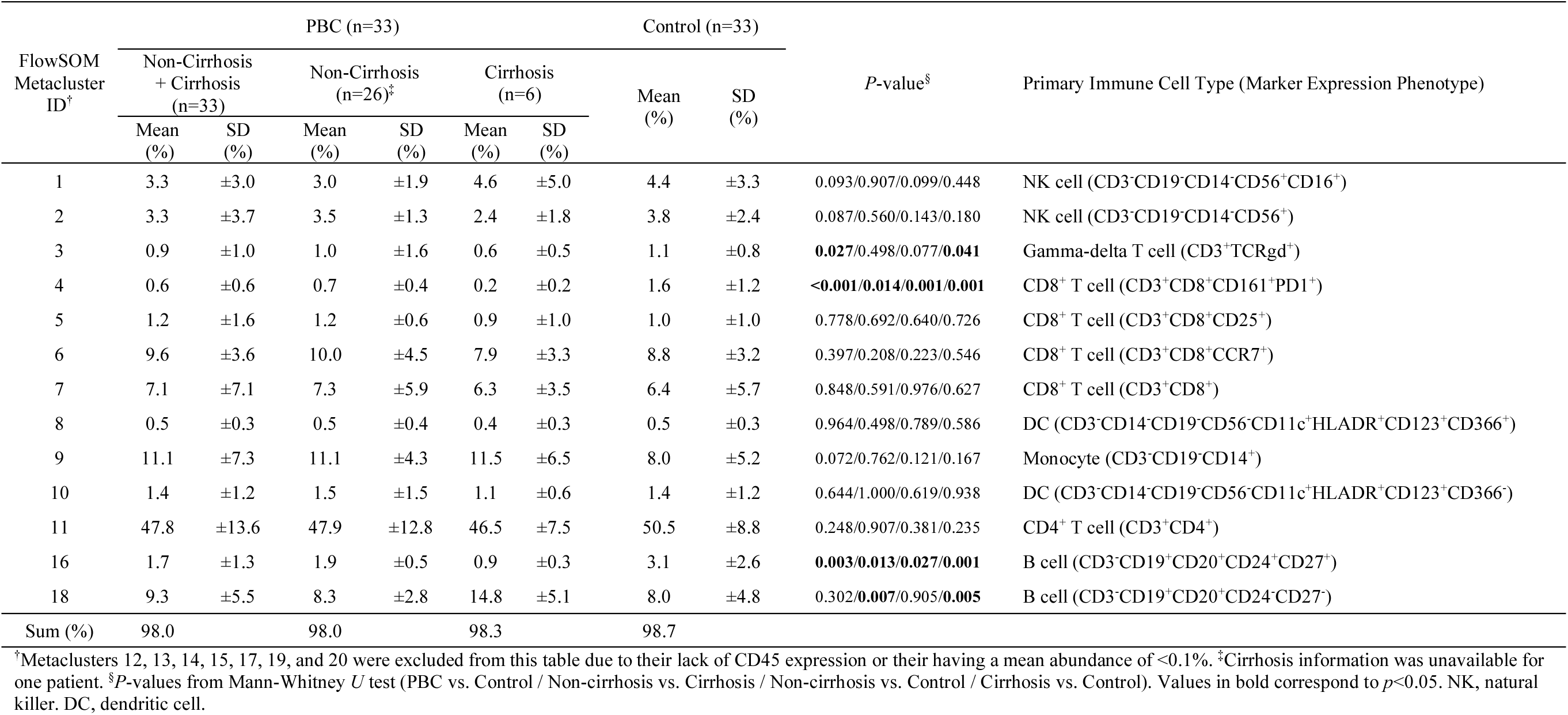
FlowSOM metacluster analysis.

**Figure 3.**
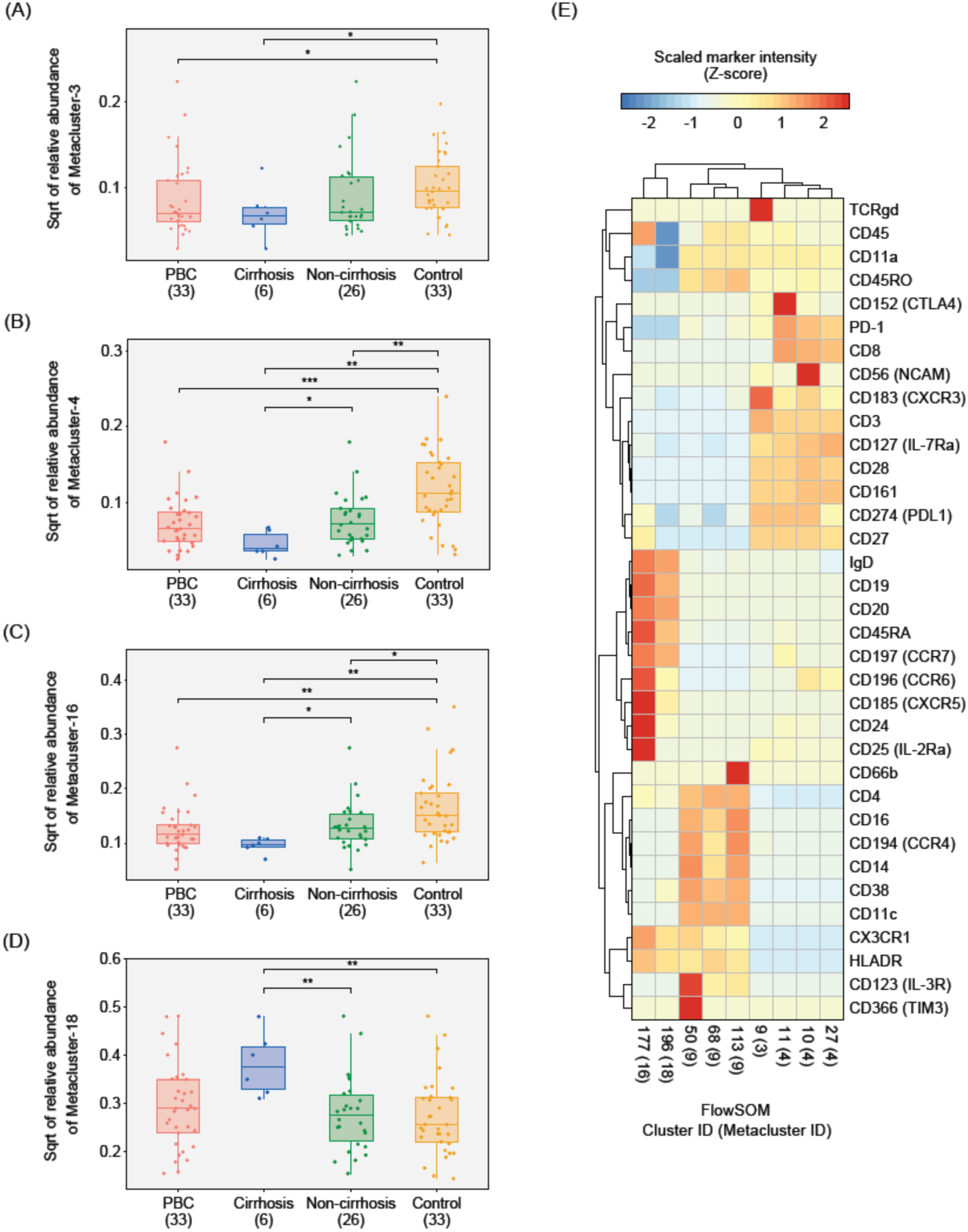
FlowSOM metaclusters, which represent immune cell types, characterize differences in immune profiles between PBC and control. (A–D) FlowSOM metaclusters that varied in relative abundance, i.e., proportion, across study groups. Boxplots indicate relative abundances of Metacluster-3, Metacluster-4, Metacluster-16, and Metacluster-18, which corresponds to gamma-delta T cell (CD3^+^TCRgd^+^), CD8^+^ T cell (CD3^+^CD8^+^CD161^+^PD1^+^), memory B cell (CD3^-^CD19^+^CD27^+^CD38^-^), and naïve B cell (CD19^+^CD27^-^IgD^+^CCR7^+^) immune cell types, respectively. The ‘Cirrhosis’ and ‘Non-cirrhosis’ groups are subsets of the ‘PBC’ group. Numbers in parentheses indicate sample size of the group. Cirrhosis status was unavailable for one PBC patient. Horizontal bars indicate the Mann-Whitney *U* test performed on the respective group pairs (significance level: *0.01≤*p*<0.05; **0.001≤*p*<0.01; and \*\*\**p*<0.001). (E) Marker expression patterns for each of the nine FlowSOM clusters (derived from Metaclusters-3, −4, −16, and −18) found to be differentially abundant (i.e., fold-change≥2.0 and *p*<0.05) between PBC and control. Clusters from the same metacluster grouped according to their marker intensities.

### Marker expression patterns reveal immune cell subsets associated with PBC

Metacluster-level analysis provides an important means to safe-guard against the potential of FlowSOM to over-fractionate cells into multiple clusters with similar marker expression profiles, which could diminish our ability to detect significant differences. However, the metacluster approach also has the potential to over-combine clusters, as evidenced by our Metacluster-11, which apparently contains all of the CD4^+^ T cells (**Table 2**). As the markers included in our panel could facilitate further characterization of these metaclusters, we also considered the association of individual immune cell clusters with PBC. Based on our statistical analysis (i.e. significant difference in abundances consisting of a minimum fold-change of 2 and *p* < 0.05), nine (among a total of 256) FlowSOM clusters were considered to be associated with PBC (**Table 3** and **Supplementary Fig. S3**; marker expression for these clusters are shown in the heatmap in **Fig. 3E**). Five of these clusters are members of the aforementioned PBC-associated metaclusters; these include: **(i)** Cluster-9 (Metacluster-3), a subset of gamma-delta T cells expressing CXCR3; **(ii)** Cluster-10 (Metacluster-4), a subset of CD161^+^CD8^+^ T cells expressing high levels of CD56; **(iii)** Cluster-11 (Metacluster-4), a subset of CD161^+^CD8^+^ T cells expressing high levels of CTLA4; **(iv)** Cluster-27 (Metacluster-4), a subset of CD161^+^PD1^+^CD8^+^ T cells; and **(v)** Cluster-177 (Metacluster-16), a subset of B cells expressing high levels of CD24 and CX3CR1. Four of the PBC-associated clusters were not members of PBC-associated metaclusters; these include: **(i)** Cluster-50 (Metacluster-9), a subset of CD14^+^CD16^+^ monocytes expressing high levels of IL3R and TIM3; **(ii)** Cluster-68 (Metacluster-9), a subset of monocytes with low CD14 expression; **(iii)** Cluster-113 (Metacluster-9), a subset of CD66b^+^ cells (contaminating granulocytes); and **(iv)** Cluster-196 (Metacluster-18), a subset of CD45^-^ cells expressing B cell markers (CD19 and CD20).

**Table 3.** FlowSOM clusters differentially abundant between PBC and Control.

| FlowSOM<br>Cluster<br>ID | FlowSOM<br>Metacluster<br>ID | Immune Cell Subset | PBC |  | Control |  | Fold-change<sup>†</sup> | *P*-value<sup>‡</sup> |
| --- | --- | --- | --- | --- | --- | --- | --- | --- |
|  |  |  | Mean<br>(%) | SD<br>(%) | Mean<br>(%) | SD<br>(%) |  |  |
| 9 | 3 | Gamma-delta T cells (CXCR3<sup>+</sup>) | 0.19 | ±0.32 | 0.37 | ±0.47 | 2.0(-) | 0.014 |
| 10 | 4 | CD8<sup>+</sup> T cells (CD161<sup>+</sup>PD1<sup>+</sup>CD56<sup>+</sup>) | 0.16 | ±0.21 | 0.56 | ±0.64 | 3.5(-) | 0.003 |
| 11 | 4 | CD8<sup>+</sup> T cells (CD161<sup>+</sup>PD1<sup>+</sup>CTLA4<sup>+</sup>) | 0.16 | ±0.15 | 0.32 | ±0.28 | 2.0(-) | 0.002 |
| 27 | 4 | CD8<sup>+</sup> T cells (CD161<sup>+</sup>PD1<sup>+</sup>) | 0.31 | ±0.38 | 0.75 | ±0.57 | 2.4(-) | <0.001 |
| 50 | 9 | Monocytes (IL3R<sup>+</sup>TIM3<sup>+</sup>) | 0.52 | ±0.60 | 0.26 | ±0.45 | 2.0(+) | 0.032 |
| 68 | 9 | Monocytes | 0.63 | ±1.08 | 0.26 | ±0.42 | 2.4(+) | 0.035 |
| 113 | 9 | CD66b<sup>+</sup> Granulocytes (contaminant) | 0.90 | ±0.84 | 0.37 | ±0.50 | 2.4(+) | 0.006 |
| 177 | 16 | Memory B cells | 0.25 | ±0.29 | 0.71 | ±0.25 | 2.9(-) | 0.004 |
| 196 | 18 | CD45<sup>-</sup> cells | 0.59 | ±0.60 | 0.26 | ±0.25 | 2.3(+) | 0.007 |

<sup>†</sup>Ratio of larger mean to smaller mean. Positive and negative signs indicate higher mean in PBC and in Control, respectively. <sup>‡</sup>*P*-values from Mann-Whitney *U* test. FlowSOM clusters were considered to be associated with PBC if fold-change was $\geq 2.0$ and $p < 0.05$ .

### Naïve B cell subsets differ in abundance between PBC patients with and without liver cirrhosis

We next explored links between disease severity and immune cell-type abundance in PBC by applying the same FlowSOM analysis to compare PBC patients with liver cirrhosis to those without (**Table 2** and **Fig. 3B–D**). Metacluster-4 (**Fig. 3B**) and Metacluster-16 (**Fig. 3C**), which were shown to be of reduced abundance in PBC patients compared to controls, also demonstrated reduced abundance in cirrhotic relative to non-cirrhotic PBC patients (3.5 fold and 2.1 fold reductions, respectively). Metacluster-18 (**Fig. 3D**) was found to have increased relative abundance in cirrhotic PBC patients compared to controls (14.8% vs. 8.0%; *p*<0.01) and to non-cirrhotic PBC patients (14.8% vs. 8.3%; *p*<0.01). This metacluster, which was found to be comprised of twenty individual clusters, expresses markers consistent with a non-specific B cell population (CD3^-^CD19^+^CD20^+^). In this regard, to better define the B cell subtype(s) responsible for the Metacluster-18 association, we identified which of the constituent clusters were individually associated with cirrhosis in PBC patients. Five of the twenty clusters of Metacluster-18 were found to be associated with cirrhosis, all of which were increased in cirrhotic PBC patients compared to non-cirrhotic PBC patients (**Table 4**). All of these clusters express markers consistent with naïve B cells (CD19^+^CD27^-^IgD^+^CCR7^+^) with minor variability in expression of chemokine receptors and other markers.

**Table 4.**
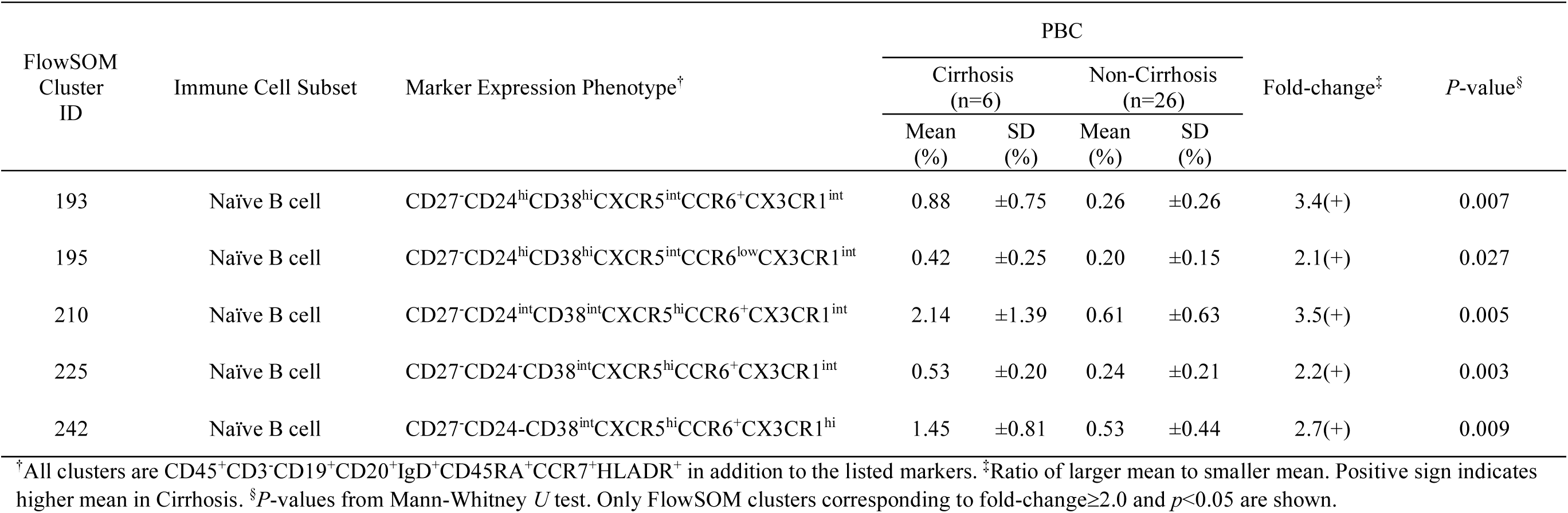
Marker expression phenotypes and abundances of cirrhosis-associated clusters belonging to Metacluster-18.

### CITRUS confirms immune cell subsets previously identified as differentially abundant in PBC

As a complement to our FlowSOM analyses, we used CITRUS to analyze the immune cell profiles of PBC patients and controls. CITRUS identified a total of 162 clusters (**Fig. 4A**), of which three were found to be differentially abundant between the PBC patients and controls. CITRUS maps overlaid with marker intensities (**Fig. 4B–D**) identify the clusters as: **(i)** a subset of B cells expressing CD27 and a high level of CD24 (**Fig. 4B**); **(ii)** a subset of CD66b^+^ cells (contaminating granulocytes) (**Fig. 4C**); and **(iii)** a subset of CD8^+^ T cells expressing high levels of CD161 (**Fig. 4D**). Notably, these results are consistent with PBC-associated clusters from the FlowSOM analysis.

**Figure 4.**
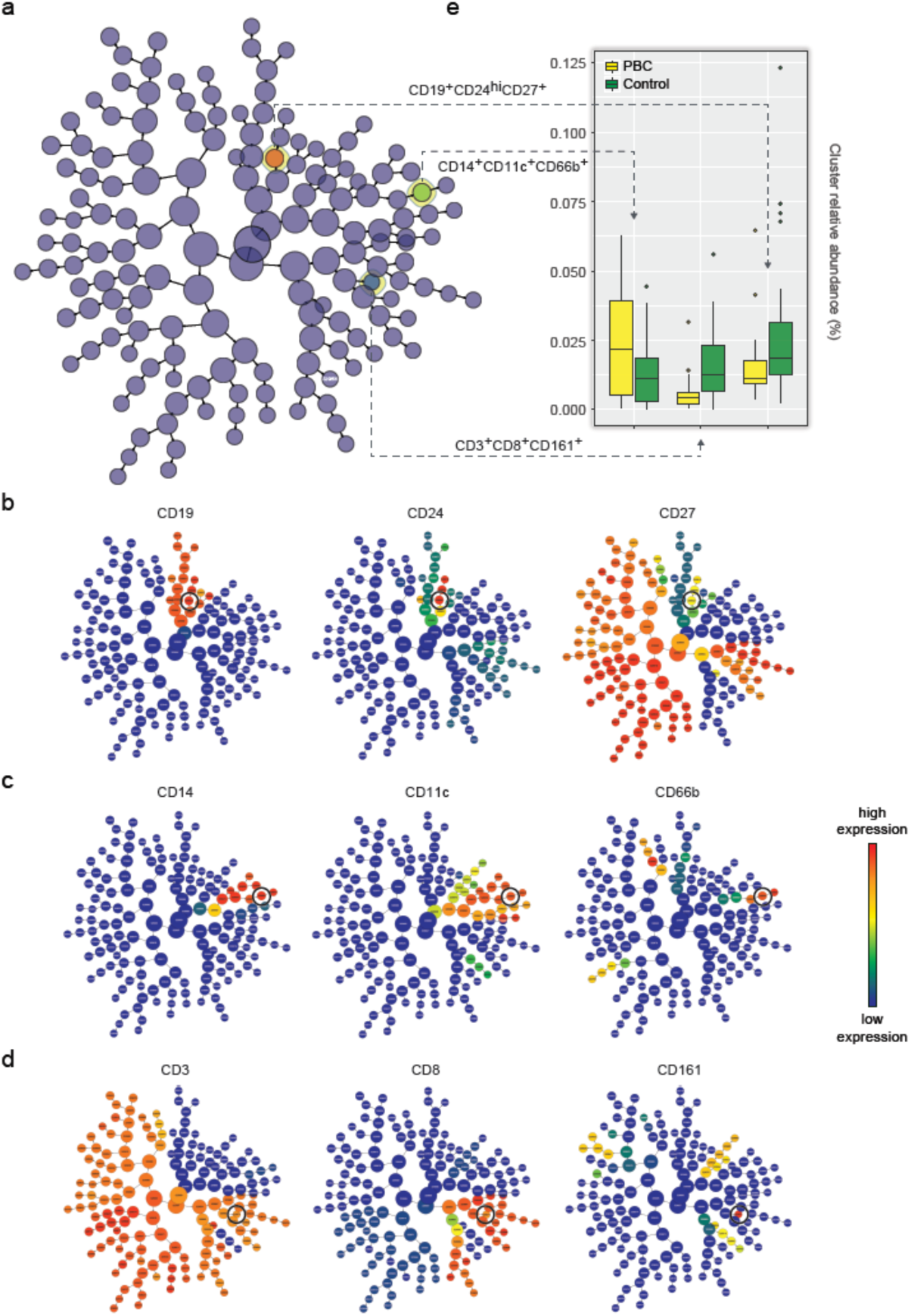
CITRUS identifies differentially abundant immune cell subsets between PBC patients and age-/sex-matched controls. (A) CITRUS produces a radial hierarchical tree of cells subsets using an unsupervised clustering approach. Each node (i.e., cluster) represents a subset of cells, and each edge points from a parent node to child node(s). Only the highlighted nodes correspond to cell subsets of statistically significant differential abundance using a significance analysis of microarray (SAM) correlative association model (Benjamini-Hochberg adjusted *P*-value<0.05). CITRUS maps overlaid with marker-specific intensities show their relative expression levels (across all nodes) in the proposed (B) CD19^+^CD24^hi^CD27^+^; (C) CD14^+^CD11c^+^CD66b^+^; and (D) CD3^+^CD8^+^CD161^+^ cell subsets. The nodes circled in black indicate positive or high expression of a particular marker. (E) CITRUS map shows a higher abundance of CD14^+^CD11c^+^CD66b^+^ cells, and lower abundances of CD19^+^CD24^hi^CD27^+^ and CD3^+^CD8^+^CD161^+^ cells, in PBC compared to control.

### Neural network classification model to distinguish PBC from control

Finally, we used our FlowSOM data set to construct a neural network model capable of discriminating PBC patients from controls based on immune profiles alone. Performance evaluation using ten-fold cross-validation demonstrated that our classification model achieved reasonably high accuracy: area under the ROC curve of 0.86 (**Supplementary Fig. S4A**); an overall classification accuracy of 86.4% (57/66) with PBC- and control-specific classification accuracies of 90.9% (30/33) and 81.8% (27/33), respectively (**Supplementary Fig. S4B**).

## Discussion

Our study is the first to perform mass cytometry-based immunophenotyping to identify distinct immune cell subsets associated with PBC. Better understanding of how the immune system contributes to, and is affected by, the underlying liver damage and resulting cholestasis in PBC could provide novel insights leading to improved prognosis, management, and therapy of this complex, multifactorial disease. While liver tissue is not amenable to regular sampling, the peripheral immune system is easily accessable, previously shown to be altered in PBC, and well-suited to routine measurement. Recent advances in flow- and mass-cytometry have greatly increased the number of cellular markers that can be simultaneously detected, allowing for a more comprehensive assessment of the immune system at single-cell resolution.

Using viSNE and FlowSOM-generated MST maps, we found that abundances and substructures of T cell, B cell, NK cell, and monocyte populations visually appear to differ between PBC patients and controls. Using FlowSOM clustering analysis, we demonstrated that metaclusters containing gamma-delta T cells, CD8^+^ T cells expressing CD161, and memory B cells expressing CD24 are of lower abundance in PBC patients relative to controls. Refined cluster analysis highlights cell subsets within these lower-abundant metaclusters, including gamma-delta T cells expressing high levels of CXCR3; CD8^+^CD161^+^ T cells expressing high levels of CD56; and CD8^+^CD161^+^ T cells expressing high levels of CTLA4. Cluster analysis also identified an interesting population of CD14^+^CD16^+^ monocytes expressing high levels of IL3R and TIM3, which were found to have significantly elevated abundance in PBC patients relative to controls. Notably, additional analysis using CITRUS was consistent with the FlowSOM findings. FlowSOM analysis also showed that two of the PBC-associated metaclusters, i.e. those containing CD8^+^CD161^+^ T cells and CD24 expressing memory B cells, were also of significantly lower abundance in cirrhotic compared to non-cirrhotic PBC patients. Moreover, this analysis also identified a non PBC-associated metacluster containing a number of naïve B cell subsets, which was found to be more abundant in cirrhotic compared to non-cirrhotic patients. Finally, we trained a neural network algorithm to differentiate PBC patients from controls, which demonstrated ∼86% accuracy in cross-validation.

Identication of differentially abundant immune cell subsets offers a glimpse into potential immune mechanisms contributing to PBC. Gamma-delta T cells are non-conventional T cells with a restricted T cell receptor repertoire often conceived as forming a bridge between innate and adaptive immunity^20^. In our study, the peripheral gamma-delta T cell population was found to be reduced in PBC compared to controls, visually on the FlowSOM-MST map and quantitatively in FlowSOM metacluster and cluster analyses. This finding is consistent with a previous study of PBC patients^21^, but at odds with other studies that reported no significant difference in gamma-delta T cell abundance between PBC patients and controls^22,23^. One of the latter studies did report an elevation of Vdelta1 gamma-delta T cells, which is a subset that is generally resident in the intestine and liver, in PBC patients with active disease or those who poorly respond to UCDA treatment relative to controls and adequate responders, respectively. Specifically targeting this subset with additional markers will be of interest in our future efforts.

CD8^+^ T cells expressing high levels of the C-type lectin CD161 include a population of unconventional innate-like T cells known as MAIT cells^24^. In our study, we found a metacluster and three of its constituent clusters expressing markers consistent with this population, and also to have significantly reduced abundance in PBC patients compared to controls. This finding is consistent with previous studies that reported reduced abundance of MAIT cells in peripheral blood^10,25^ and in liver tissues^10^ of PBC patients compared to healthy controls. Setsu *et al*. observed that depleted MAIT cells were not recovered to normal levels in the blood of PBC patients even after UDCA treatment, suggesting the possibility that MAIT cells were being activated, exhausted, and depleted due to ongoing liver inflammation^10^. Along those lines, the reduced-abundance CD8^+^CD161^+^ T cell metacluster in our study also expressed high levels of the immune-exhaustion biomarker PD-1, and one of the constituent clusters expressed high-levels of CTLA4, which is also a marker of exhaustion. Moreover, the abundance of this metacluster was reduced even further in cirrhotic compared to non-cirrhotic PBC patients. This finding is consistent with studies of other chronic liver diseases, including hepatitis B viral infection^26^ and hepatocellular carcinoma^27^, which implicate exhaustion and loss of MAIT cells with worsening liver damage. Considering the apparent importance of these cells in PBC, our future efforts should include additional MAIT-specific markers such as the MAIT-invariant T cell receptor (Vα7.2).

The CD24^+^CD27^+^ B cell population contains subsets of regulatory B cells (B_reg_) that include plasmablasts, transitional B cells and B10 cells, which are important contributers to immune homeostasis and maintenance of tolerance^28^. In our study, abundance of a metacluster and one of its constituent clusters expressing markers consistent with this population was found to be markedly decreased in PBC patients relative to controls, and in cirrhotic compared to non-cirrhotic PBC patients. This finding is in contrast to other studies in PBC, which identified no such differences^29,30^. However, the approaches in those studies were quite different from ours, so the disparity should be interpreted with caution. Among the B_reg_ subtypes, the PBC- and cirrhosis-associated cluster most closely represented B10 cells, due to the lack of CD38 and high expression of IgD^31^. This rare subtype of B cells mediate immune responses through their secretion of anti-inflammatory cytokine IL-10, and have been implicated in autoimmunity^32^. Considering our findings, along with the known anti-inflammatory effects of B10 cells and their therapeutic potential^33^, more in-depth study of the role of memory and regulatory B cells, particularly B10 cells, in PBC is strongly warranted.

Three clusters belonging to the metacluster containing monocytes were found to have increased abundance in PBC relative to controls. One of the groups strongly expressed CD66b and was considered to represent granulocyte contamination in the PBMC fraction. The second cluster expressed CD16 and low level of CD14, consistent with the classification of non-classical monocyte. While these cells are generally considered to be involved with resolving inflammation, some studies suggest they may be pathogenic in the context of certain inflammatory diseases^34^. The third cluster expressed CD14 and CD16, consistent with the classification of intermediate monocyte, but also expressed high levels of the IL3 receptor CD123, and the immunomodulatory molecule TIM3. While both CD123 and TIM3 have been shown to modulate the function of monocytes and other innate immune cells^35^, there is currently no evidence on how this rare population of cells could contribute to PBC. Monocytes and other myeloid cells present in the PBMC fraction share many of the same markers; however, as our CyTOF immunophenotyping panel did not include many myeloid-specific markers, it is difficult to discriminate between these subtypes in the current study. Future efforts in PBC should consider inclusion of a more diverse set of markers to assist in characterizing myeloid subtypes^36^.

B cells are widely acknowledged to contribute to the immunopathogenesis and maintenance of PBC^37–39^. We identified a metacluster and five of its constituent clusters, all of which expressing markers indicative of naïve B cell-like properties, to be of higher abundance in cirrhotic compared to non-cirrhotic patients. Notably, these clusters were not found to differ in the full set of PBC patients relative to controls. While this finding should be taken with caution due to the low number of cirrhotic patients in the study, it does point to the need for an expanded set of B cell-specific markers in future studies of PBC.

While small in sample size, our study was able to identify a number of immune cell subsets demonstrating altered abundance in PBC patients relative to controls, and in cirrhotic compared to non-cirrhotic PBC patients. Findings such as the reduced abundance of MAIT cells in PBC are consistent with the existing literature, suggesting that our stored PBMCs are of adequate quality to obtain meaningful results. Moreover, detection of immune cell subsets that show differential abundance in patients with more advanced disease suggests that immune profiling could eventually become a valuable tool for the clinical management of PBC. Our study warrants further efforts that include additional cell type-specific markers, a wider range of analytical modalities, and individuals representing a broader range of disease phenotypes and severities. In this regard, we expect that such future investigations will lead to a better understanding of PBC pathogenesis and potential discovery of novel biomarkers and treatment modalities that could benefit PBC patients.

## Acknowledgements

The authors would like to thank Michael A. Strausbauch, Kevin D. Pavelko, and the Mayo Clinic Immune Monitoring Core for CyTOF support, and Drs. Kara Conway and Bernard Khor for helpful discussions. This work was supported by the Mayo Clinic Center for Individualized Medicine (J.S.J, K.Y.C., V.K.G., J.S.), Mark E. and Mary A. Davis to Mayo Clinic Center for Individualized Medicine (J.S.), National Institutes of Health K12 HD065987 (E.A.L.E.), Everett J. and Jane M. Hauck Associate Director endowment for the Center for Individualized Medicine, Mayo Clinic (K.N.L.), and the William O. Lund, Jr. and Natalie C. Lund Charitable Foundation endowment for the Center for Individualized Medicine, Mayo Clinic (K.N.L.).

## Author contributions

K.N.L. conceived the study. J.S.J., B.J., and J.S. designed the analytical metholodogies. J.S.J, K.Y.C., V.K.G., and J.S. conducted the experiments. J.S.J, B.J., K.Y.C., V.K.G., Y.M.S., J.D.Y., A.H.A., E.A.L.E., J.S., and K.N.L. analyzed and interpreted the results. J.S.J., B.J., J.S., and K.N.L. wrote the manuscript, with editorial contributions from other co-authors. All authors reviewed and approved the final version of the manuscript.

## Additional information

Correspondence and requests for materials should be addressed to J.S. or K.N.L.

## Data availability

Raw mass cytometry data will be made publicly available as a report on Cytobank (http://www.cytobank.org) with linked flow cytometry standard (.fcs) files upon acceptance of the manuscript.

