## Supplementary Information for "Single-Cell Mass Cytometry on Peripheral Blood Identifies Immune Cell Subsets Associated with Primary Biliary Cholangitis"

Supplementary Table S1. List of primary antibodies used in this study.

| No. | Species | Clone | Metal | Target | Amount required per test (μL) |
|---|---|---|---|---|---|
| 1 | Human | 3G8 | 148Nd | CD16 | 0.125 |
| 2 | Human | SK1 | 168Er | CD8a | 0.125 |
| 3 | Human | 2H7 | 171Yb | CD20 | 0.125 |
| 4 | Human | SK3 | 174Yb | CD4 | 0.125 |
| 5 | Human | HI30 | 089Y | CD45 | 0.250 |
| 6 | Human | HIB19 | 142Nd | CD19 | 0.250 |
| 7 | Human | IA6-2 | 146Nd | IgD | 0.250 |
| 8 | Human | Bu15 | 147Sm | CD11c | 0.250 |
| 9 | Human | 11F2 | 152Sm | TCRgd | 0.250 |
| 10 | Human | UCHT1 | 154Sm | CD3 | 0.250 |
| 11 | Human | HI100 | 155Gd | CD45RA | 0.250 |
| 12 | Human | L128 | 158Gd | CD27 | 0.250 |
| 13 | Human | L243 | 173Yb | HLA-DR | 0.250 |
| 14 | Human | G034E3 | 141Pr | CD196 (CCR6) | 0.500 |
| 15 | Human | A019D5 | 143Nd | CD127 (IL-7R) | 0.500 |
| 16 | Human | HIT2 | 144Nd | CD38 | 0.500 |
| 17 | Human | L291h4 | 149Sm | CD194 (CCR4) | 0.500 |
| 18 | Human | 6H6 | 151Eu | CD123 (IL-3R) | 0.500 |
| 19 | Human | RF8B2 | 153Eu | CD185 (CXCR5) | 0.500 |
| 20 | Human | CD28.2 | 160Gd | CD28 | 0.500 |
| 21 | Human | G025H7 | 163Dy | CD183 (CXCR3) | 0.500 |
| 22 | Human | UCHL1 | 165Ho | CD45RO | 0.500 |
| 23 | Human | ML5 | 166Er | CD24 | 0.500 |
| 24 | Human | G043H7 | 167Er | CD197 (CCR7) | 0.500 |
| 25 | Human | 2A3 | 169Tm | CD25 (IL-2R) | 0.500 |

| | | | | | |
|---|---|---|---|---|---|
| 26 | Human | M5E2 | 175Lu | CD14 | 0.500 |
| 27 | Human | NCAM16.2 | 176Yb | CD56 (NCAM) | 0.500 |
| 28 | Human | 80H3 | 162Dy | CD66b | 1.000 |
| 29 | Human | HP-3G10 | 164Dy | CD161 | 1.000 |
| 30 | Human | 11C3C65 | 150Nd | LAG-3 | 1.000 |
| 31 | Human | 29E.2A3 | 156Gd | PD-L1 | 1.000 |
| 32 | Human | 14D3 | 170Er | CTLA-4 | 1.000 |
| 33 | Human | 2A9-1 | 172Yb | CX3CR1 | 1.000 |
| 34 | Human | HI111 | 145Nd | CD11a | 0.250 |
| 35 | Human | F38-2E2 | 159Tb | Tim-3 | 1.000 |
| 36 | Human | EH12.2H7 | 161Dy | PD-1 | 1.000 |

Supplementary Table S2. Distribution of cell counts from all 66 PBC and control samples across three different stages of the mass cytometry experiment.

| Group | Post-thaw cell count | Viability (%) | Post-deconvolution event count | Post-QC event Count |
|---|---|---|---|---|
| PBC_1 | 23,031,054 | 1.2 | 202,003 | 65,733 |
| PBC_2 | 6,773,839 | 76.2 | 186,356 | 55,738 |
| PBC_3 | 7,436,561 | 83.7 | 456,148 | 246,364 |
| PBC_4 | 11,107,924 | 77.2 | 132,597 | 66,214 |
| PBC_5 | 4,961,617 | 88.2 | 294,370 | 215,064 |
| PBC_6 | 3,847,306 | 82.6 | 168,051 | 105,193 |
| PBC_7 | 4,070,168 | 85.0 | 144,518 | 87,427 |
| PBC_8 | 7,890,000 | 77.0 | 195,881 | 115,118 |
| PBC_9 | 6,380,000 | 93.0 | 411,839 | 290,066 |
| PBC_10 | 9,850,000 | 72.7 | 142,290 | 48,431 |
| PBC_11 | 5,330,000 | 91.7 | 273,194 | 167,477 |
| PBC_12 | 5,410,000 | 87.6 | 234,168 | 159,360 |
| PBC_13 | 7,210,000 | 79.9 | 175,132 | 115,964 |
| PBC_14 | 6,590,000 | 82.2 | 115,846 | 60,835 |
| PBC_15 | 5,860,000 | 85.2 | 48,225 | 29,516 |
| PBC_16 | 1,520,000 | 88.5 | 33,315 | 17,639 |
| PBC_17 | 9,160,000 | 75.8 | 69,586 | 39,353 |
| PBC_18 | 1,580,000 | 78.1 | 35,608 | 18,814 |
| PBC_19 | 6,000,000 | 77.7 | 91,287 | 41,036 |
| PBC_20 | 8,730,000 | 81.8 | 33,363 | 17,665 |
| PBC_21 | 5,600,880 | 80.8 | 599,286 | 397,133 |
| PBC_22 | 3,589,255 | 82.8 | 292,122 | 178,702 |
| PBC_23 | 6,474,735 | 81.5 | 353,941 | 249,787 |
| PBC_24 | 14,568,153 | 4.7 | 103,234 | 53,999 |
| PBC_25 | 3,190,449 | 78.1 | 394,524 | 238,170 |
| PBC_26 | 7,213,699 | 81.2 | 422,794 | 299,032 |

| | | | | |
|---|---|---|---|---|
| PBC_27 | 8,767,870 | 78.1 | 367,743 | 244,045 |
| PBC_28 | 2,228,623 | 94.7 | 281,392 | 156,677 |
| PBC_29 | 4,504,163 | 81.1 | 661,733 | 430,210 |
| PBC_30 | 2,797,508 | 88.1 | 515,905 | 340,659 |
| PBC_31 | 6,545,112 | 71.8 | 246,500 | 127,622 |
| PBC_32 | 7,207,834 | 74.6 | 366,443 | 218,469 |
| PBC_33 | 16,175,108 | 87.3 | 543,394 | 401,510 |
| Control_1 | 13,377,600 | 72.9 | 125,621 | 33,100 |
| Control_2 | 5,935,174 | 86.2 | 523,610 | 189,651 |
| Control_3 | 8,879,301 | 74.3 | 285,820 | 134,048 |
| Control_4 | 10,263,393 | 71.1 | 90,463 | 29,730 |
| Control_5 | 4,433,786 | 80.8 | 157,192 | 95,262 |
| Control_6 | 5,606,745 | 81.9 | 262,079 | 171,887 |
| Control_7 | 7,348,589 | 74.7 | 127,357 | 56,332 |
| Control_8 | 11,700,000 | 73.8 | 183,802 | 76,000 |
| Control_9 | 4,450,000 | 82.1 | 352,184 | 184,687 |
| Control_10 | 11,700,000 | 76.7 | 176,555 | 84,522 |
| Control_11 | 1,960,000 | 92.8 | 187,868 | 105,938 |
| Control_12 | 10,300,000 | 55.2 | 279,858 | 54,553 |
| Control_13 | 6,840,000 | 72.1 | 141,117 | 83,531 |
| Control_14 | 5,750,000 | 87.3 | 66,461 | 33,955 |
| Control_15 | 9,880,000 | 65.0 | 83,576 | 34,763 |
| Control_16 | 4,250,000 | 86.6 | 258,465 | 178,920 |
| Control_17 | 1,920,000 | 89.9 | 109,464 | 70,916 |
| Control_18 | 5,230,000 | 84.9 | 136,440 | 90,915 |
| Control_19 | 7,880,000 | 85.8 | 47,925 | 25,753 |
| Control_20 | 5,120,000 | 87.2 | 111,198 | 71,984 |
| Control_21 | 1,735,980 | 94.9 | 231,317 | 143,649 |
| Control_22 | 3,612,714 | 96.8 | 847,255 | 510,051 |
| Control_23 | 7,841,232 | 82.2 | 523,957 | 381,778 |
| Control_24 | 6,744,515 | 62.7 | 213,287 | 117,966 |

| | | | | |
|---|---|---|---|---|
| Control_25 | 4,111,222 | 96.4 | 674,103 | 485,811 |
| Control_26 | 4,240,248 | 95.6 | 644,313 | 477,152 |
| Control_27 | 5,624,339 | 88.6 | 509,045 | 346,213 |
| Control_28 | 2,867,885 | 90.4 | 338,518 | 212,315 |
| Control_29 | 4,744,620 | 84.5 | 351,794 | 187,192 |
| Control_30 | 4,375,138 | 79.6 | 640,654 | 445,053 |
| Control_31 | 6,433,681 | 79.9 | 428,364 | 299,692 |
| Control_32 | 6,052,470 | 93.8 | 801,474 | 477,574 |
| Control_33 | 6,087,658 | 77.9 | 338,325 | 216,743 |

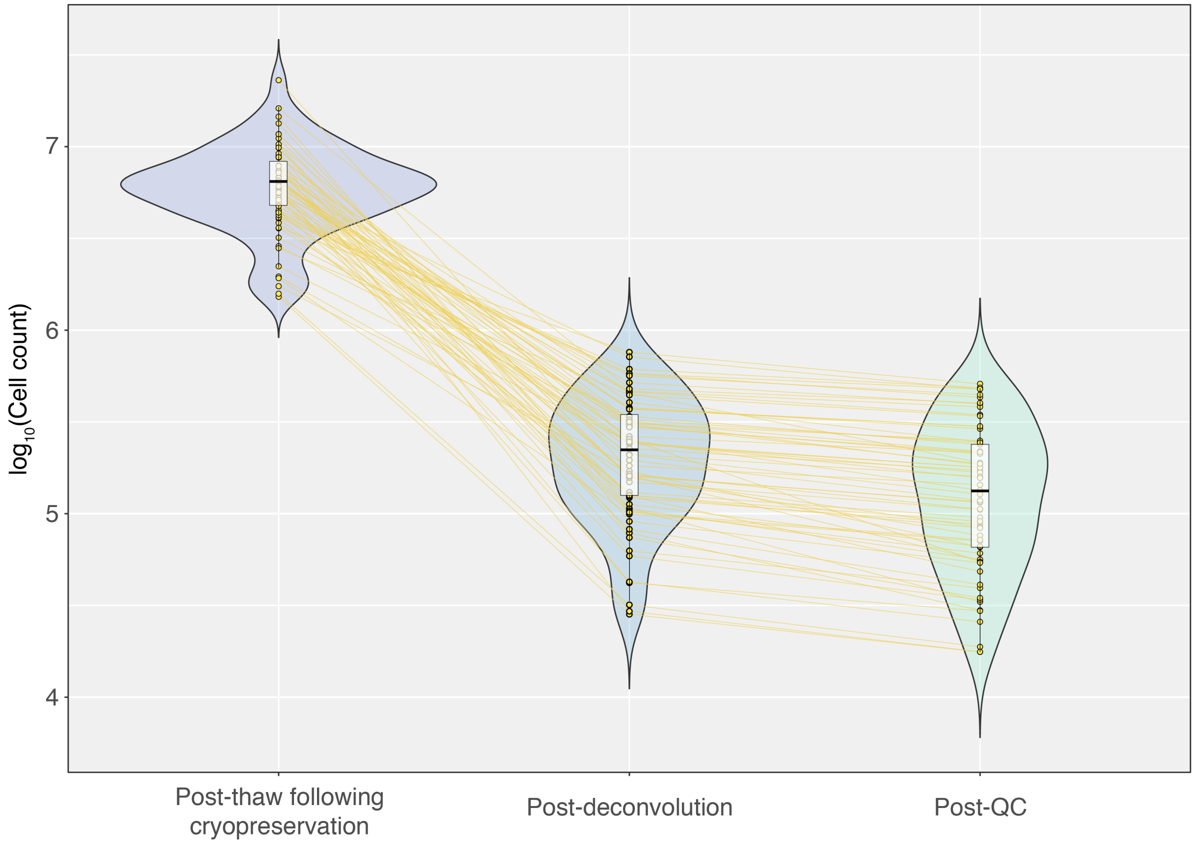


**Supplementary Figure S1.** Distribution of cell counts from all 66 PBC and control samples across three different stages of the mass cytometry experiment. Gold lines connect points corresponding to the same sample. Post-thaw following cryopreservation: The number of viable and non-viable cells counted after thawing following cryopreservation (or the original number of cells that were cyropreserved). These numbers corresponds to the total yield of cells before metal-conjugated antibody labeling; Post-deconvolution: The number of events in each sample after antibody labeling and sample debarcoding (i.e., deconvolution); Post-QC: The number of events in each sample after the quality-control process (which includes elimination of debris, dead cells, and doublets) indicating the actual number of viable cells that were profiled.

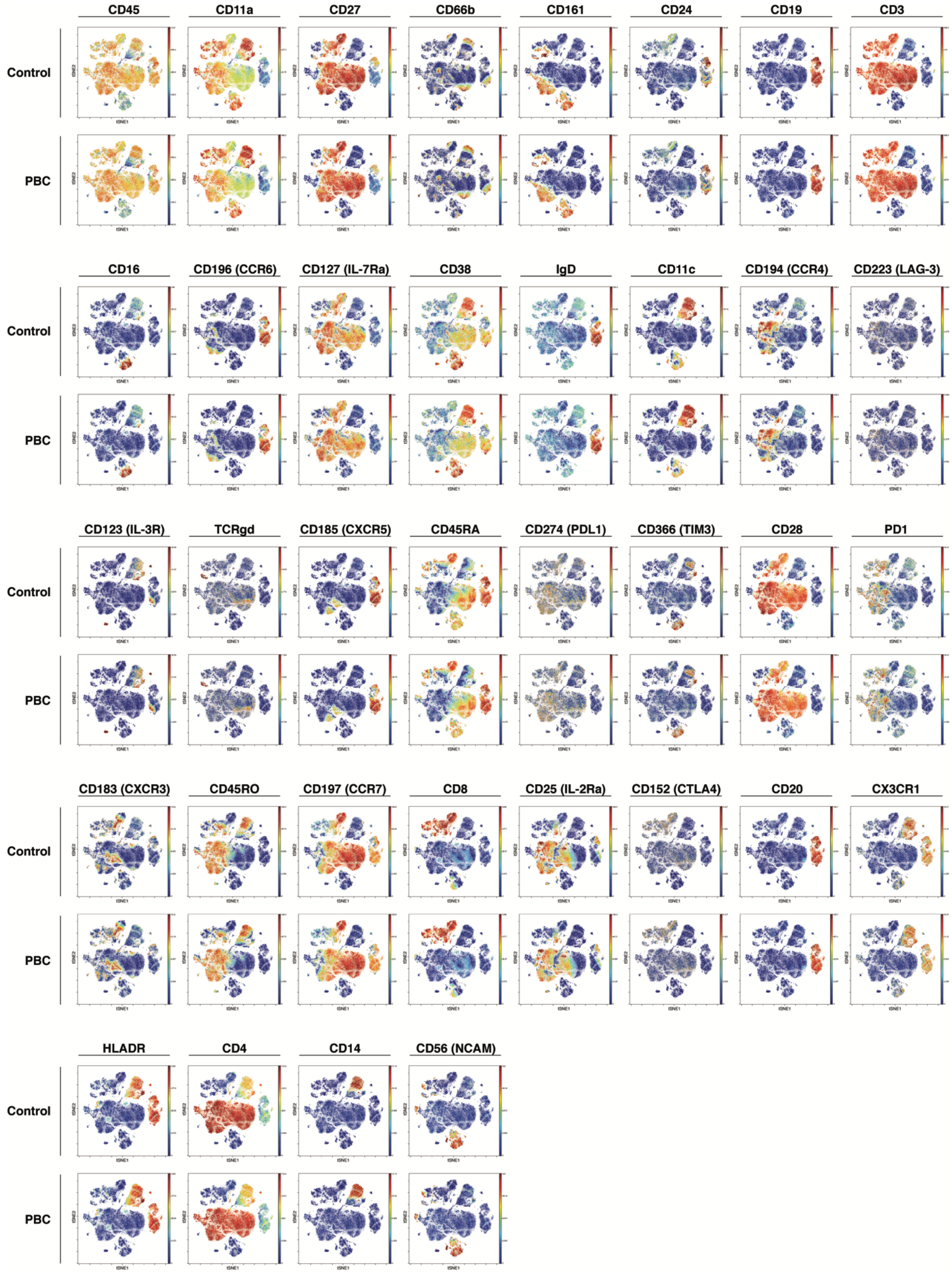
CD45
CD11a
CD27
CD66b
CD161
CD24
CD19
CD3
Control
PBC
CD16
CD196 (CCR6)
CD127 (IL-7Ra)
CD38
IgD
CD11c
CD194 (CCR4)
CD223 (LAG-3)
CD123 (IL-3R)
TCRgd
CD185 (CXCR5)
CD45RA
CD274 (PDL1)
CD366 (TIM3)
CD28
PD1
CD183 (CXCR3)
CD45RO
CD197 (CCR7)
CD8
CD25 (IL-2Ra)
CD152 (CTLA4)
CD20
CX3CR1
HLADR
CD4
CD14
CD56 (NCAM)
tSNE1
tSNE2

**Supplementary Figure S2.** t-SNE maps colored by channel intensities show relative expression of immune cell markers. Mass cytometry samples from 33 PBC patients were concatenated into 100,000 randomly-sampled total events and mapped onto a t-SNE plot using viSNE (PBC). Analogously, samples from 33 age-/sex-matched controls were mapped using viSNE (Control). Each point in the t-SNE plot represents a single event (e.g., cell) detected by the mass cytometer, and colors vary according to the degree of marker expression. Observed regional differences in the viSNE maps correspond to differences in expression patterns of immune cell markers.

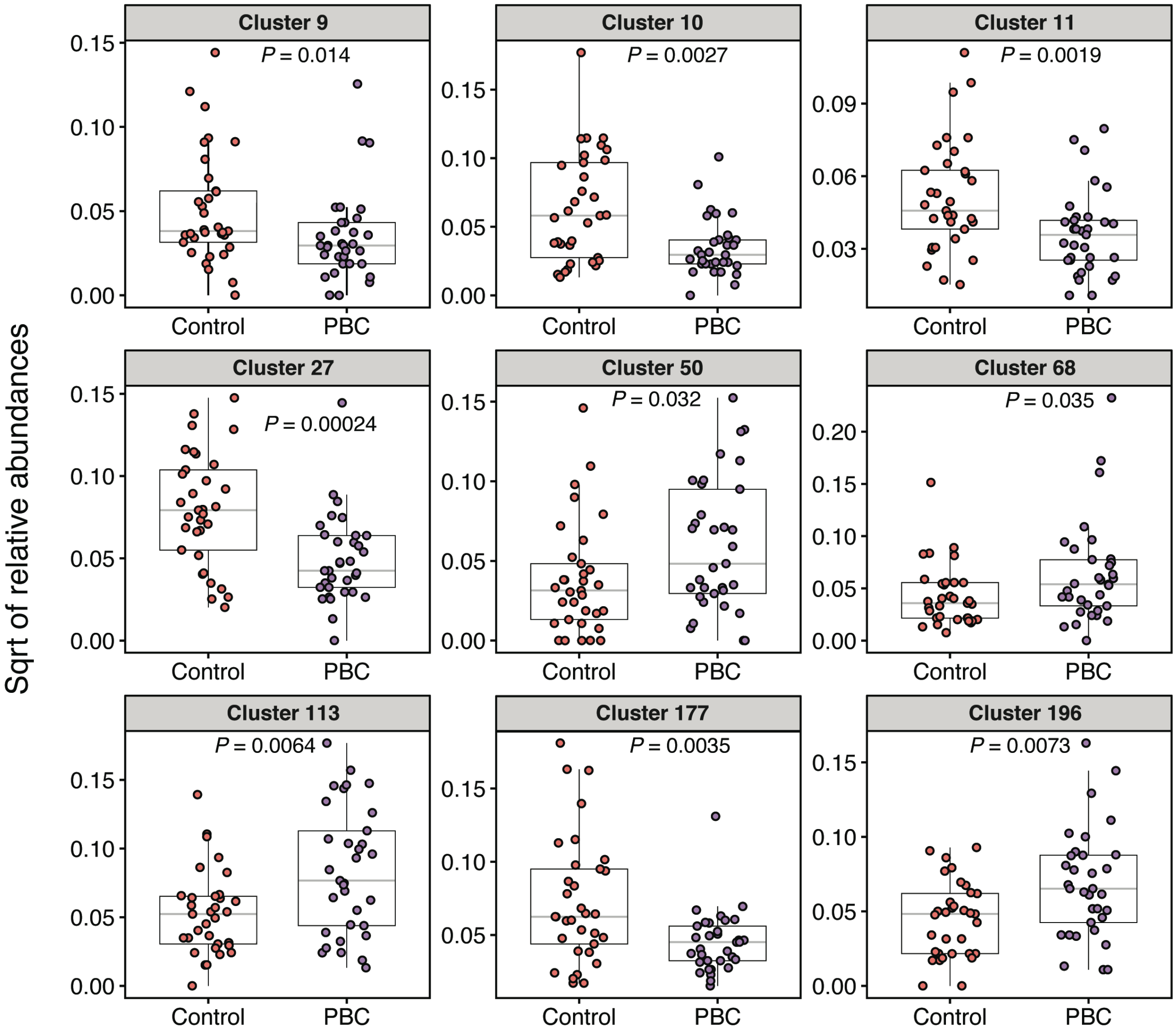


**Supplementary Figure S3.** FlowSOM clusters whose relative abundances were significantly different between PBC and control (fold-change≥2.0 and $p$<0.05 based on Mann-Whitney $U$ test).

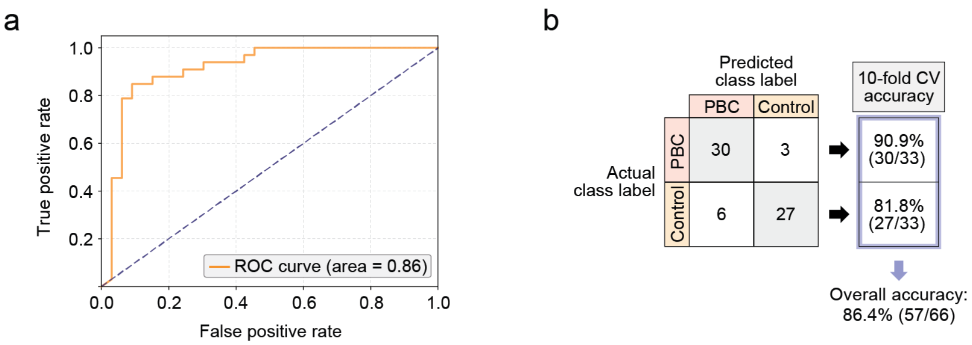


**Supplementary Figure S4.** A neural network-based classification model, which was trained upon FlowSOM cluster features and class labels, was able to distinguish immune profiles of PBC from those of control with high prediction accuracy. Model performance was evaluated in ten-fold cross-validation. (a) ROC curve with an AUC of 0.86; and (b) confusion matrix of model predictions, demonstrating an overall phenotype prediction accuracy of 86.4% and group-wise accuracies of 90.9% and 81.8% for PBC and control, respectively.
